# Open Data Commons for Preclinical Traumatic Brain Injury Research: Empowering Data Sharing and Big Data Analytics

**DOI:** 10.1101/2021.03.15.435178

**Authors:** Austin Chou, Abel Torres Espin, J.Russell Huie, Karen Krukowski, Sangmi Lee, Amber Nolan, Caroline Guglielmetti, Bridget E. Hawkins, Myriam M. Chaumeil, Geoffrey T. Manley, Michael S. Beattie, Jacqueline C. Bresnahan, Maryann E. Martone, Jeffrey S. Grethe, Susanna Rosi, Adam R. Ferguson

**Author notes:** Corresponding Authors: Adam R Ferguson, Ph.D., Associate Professor, Zuckerberg San Francisco General Hospital, Building #1 Room 101, San Francisco, CA 94110, Susanna Rosi, Ph.D., Lewis and Ruth Cozen Chair II, Professor, 1001 Potrero Ave, Zuckerberg San Francisco General Hospital, Building #1 Room 101, San Francisco, CA 94110.

## Abstract

Traumatic brain injury (TBI) is a major unsolved public health problem worldwide with considerable preclinical research dedicated to recapitulating clinical TBI, deciphering the underlying pathophysiology, and developing therapeutics. However, the heterogeneity of clinical TBI and correspondingly in preclinical studies have made translation from bench to bedside difficult. Here, we present the potential of data sharing, data aggregation, and multivariate analytics to integrate heterogeneity and empower researchers. We introduce the Open Data Commons for Traumatic Brain Injury (ODC-TBI.org) as a user-centered web platform and cloud-based repository focused on preclinical TBI research that enables data citation with persistent identifiers, promotes data element harmonization, and follows FAIR data sharing principles. Importantly, the ODC-TBI implements data sharing at the level of individual subjects, thus enabling data reuse for granular big data analytics and data-hungry machine learning approaches. We provide use cases applying descriptive analytics and unsupervised machine learning on pooled ODC-TBI data. Descriptive statistics included subject-level data for 11 published papers (N = 1250 subjects) representing six distinct TBI models across mice and rats (implementing controlled cortical impact, closed head injury, fluid percussion injury, and CHIMERA TBI modalities). We performed principal component analysis (PCA) on cohorts of animals combined through the ODC-TBI to identify persistent inflammatory patterns across different experimental designs. Our workflow ultimately improved the sensitivity of our analyses in uncovering patterns of pro- vs anti-inflammation and oxidative stress without the multiple testing problems of univariate analyses. As the practice of open data becomes increasingly required by the scientific community, ODC-TBI provides a foundation that creates new scientific opportunities for researchers and their work, facilitates multi-dataset and multidimensional analytics, and drives collaboration across molecular and computational biologists to bridge preclinical research to the clinic.

## Introduction

Traumatic brain injury (TBI) is a leading cause of neurological disorders and affects over 69 million people annually worldwide [1,2]. The incidence of TBI is expected to rise each year, and over 3 million patients in the U.S. alone and many more globally suffer from chronic TBI-related disabilities [2–4]. Despite the abundance of preclinical TBI studies, randomized controlled clinical trials have consistently failed [5,6]. The lack of effective treatment can be broadly ascribed to the heterogeneity of injuries and the varied pathological biology captured by the broad definition of TBI: a disruption of neurological function caused by a bump, blow, or jolt to the head or penetrating head injury [7].

Biological injury responses can differ dramatically across injury sites, injury severities, and patient characteristics [8]. To capture the heterogeneity of clinical TBIs, a multitude of preclinical TBI models have been developed to isolate specific injury mechanisms [9]. While the diverse injury parameters and outcome measures employed by different experimenters do effectively recapitulate distinct aspects of clinical pathology, the breadth of preclinical models and research ultimately makes inferential insights difficult to compare across studies and translate across species. Preclinical TBI models have thus largely been treated as very distinct representations of clinical TBI, circumventing the complexity of TBI heterogeneity instead of directly addressing it. However, the wealth of data collected across TBI models presents a new opportunity for rigorous joint analyses across studies and across preclinical TBI models to directly investigate common biological features underlying heterogeneity. Indeed, there is growing interest and support for the application of Big Data frameworks and multidimensional machine learning to TBI research [10–12]. Such techniques have been recently employed with clinical data to reveal TBI pathophysiology persistent across heterogeneous patients [13,14]. While similar efforts in preclinical TBI research are still nascent, they represent a unique perspective towards unraveling common pathological mechanisms and bridging preclinical to clinical research.

A major obstacle to the Big Data approach is the under-developed and under-utilized practice of data sharing and data standardization and harmonization in the preclinical TBI field. Clinical TBI data programs, such as Transforming Research and Clinical Knowledge in TBI (TRACK-TBI) and Collaborative European NeuroTrauma Effectiveness Research in TBI (CENTER-TBI), have dramatically improved access to data and enabled multidimensional analytics in clinical research [15–18]. In contrast, most preclinical TBI data and research have been communicated and shared solely through publications without the release of the underlying data. The data of each published specimen are thus sequestered as summarized aggregates which makes individual-subject level data inaccessible for data reuse and further analytics [12,19]. Additionally, the language and terminology of collected variables can differ in name and definition between labs. The National Institute of Neurological Disorders and Stroke (NINDS) have released dictionaries of “common data elements” (CDEs), basic units of data that prescribe the data type and standardize the language for variables in an effort to improve the reproducibility of clinical and preclinical TBI research [20,21]. However, there remains an unmet need for open data infrastructures that host preclinical TBI data and for datasets to begin integrating the NINDS-defined CDEs for data sharing and reusability.

In this article, we present the Open Data Commons for TBI (ODC-TBI.org), a platform and repository for data sharing for the global preclinical TBI research community. The infrastructure is developed in collaboration with the Neuroscience Information Framework (NIF) [22]. Building upon previous work on the Open Data Commons for Spinal Cord Injury (ODC-SCI.org) [23,24], we developed the ODC-TBI for protected data sharing while upholding data stewardship principles towards making biomedical data Findable, Accessible, Interoperable, and Reusable (FAIR) [25]. To jumpstart FAIR sharing in preclinical TBI, we standardized datasets from 11 publications along NINDS-defined CDEs and uploaded them to the ODC-TBI. As a proof of concept for Big Data analytics enabled by the ODC-TBI, we aggregated data from three separate experiments uploaded to the ODC-TBI and harnessed multivariate analytics to uncover persistent patterns of the inflammatory response in the controlled cortical impact TBI mouse model. Altogether, we illustrate the infrastructure of the ODC-TBI to promote data sharing within the preclinical TBI research community and demonstrate the utility of multidataset, multidimensional analytics to uncover common TBI pathophysiology across heterogeneous experimental features.

## Results

### ODC-TBI Infrastructure for Data Sharing and Security

The purpose of the ODC-TBI is to establish an infrastructure to facilitate effective data sharing practices within the preclinical TBI research community and expand the data standardization and harmonization guidelines initiated by the NINDS [20,26]. Additionally, the ODC-TBI interface has been developed to address the concerns of preclinical TBI researchers towards data sharing practices [21] and empower the researchers through an intuitive interface. Currently, the ODC-TBI provides guidelines to help researchers format their dataset according to best practices for data interoperability [27,28] and standardize them according to FAIR principles [25] and NINDS-defined CDEs. Once prepared, datasets can be uploaded to the ODC-TBI and then further combined for Big Data analytics (**Fig 1A**).

**Figure 1.**
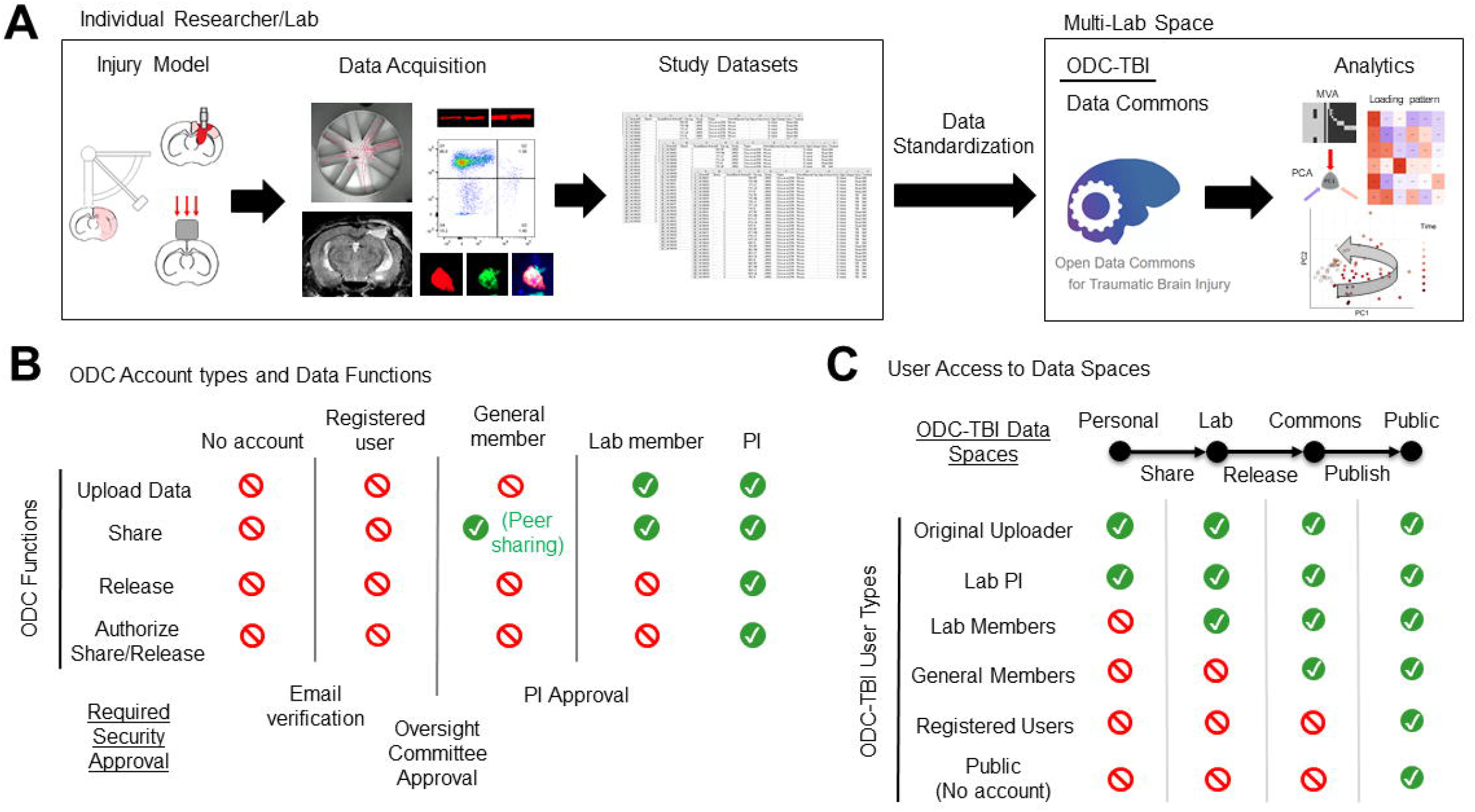
Open data commons for Traumatic Brain Injury (ODC-TBI) data flow and accessibility summary. (**A**) Experiments are currently carried out by individual researchers and labs. The resulting datasets are commonly preserved within lab resources (e.g. harddrives, lab notebooks, etc). The ODC-TBI provides documentation and guidance to help standardize datasets with respect to NINDS-defined preclinical common data elements (CDEs). Datasets from multiple labs and centers can then be uploaded into the ODC-TBI and shared and combined for further analysis. (**B**) The ODC-TBI has five user types with three steps for security. Each user type has different available functions on the site. After an email verification and approval by the ODC-TBI committee to ensure the user is a researcher in the TBI field, they become a general member. They can then join or create a lab, which requires the lab PI’s approval. Lab members can upload and share their data within the lab they have joined. Lastly, PI-level users can also initiate the dataset release/publication process, increasing the accessibility of their data to others outside of their lab. (**C**) ODC-TBI consists of four Data Spaces. Each Data Space has different levels of accessibility. Datasets are delegated to a Personal space when they are first uploaded; Personal datasets are accessible only to the uploader and their lab PI. Datasets are shared into the Lab space where they can be accessed by anyone in the lab. Datasets can be released into the Community space where other general members can access them. Lastly, PIs can publish their datasets which will make the dataset accessible to the general public as citable units of research with unique digital object identifiers.

Protecting a lab’s data from misuse by third parties is a major concern of investigators [29,30]. To address this obstacle, the ODC-TBI is built on a robust cloud-based cyberinfrastructure through the California Institute for Telecommunications and Information Technology (CalIT2) which includes e-commerce grade security and encryption. In addition, ODC-TBI has established several approval protocols to provide qualified access to sensitive data while enabling open access to published data (**Fig 1B**). Uploading, sharing, and accessing data is only possible for users who have a verified institutional email and have been approved for lab membership by a Principal Investigator (PI) with a lab in ODC-TBI. The process of data sharing requires authorization by the PI, and the PI can remove datasets from the shared space at any time. When data is first uploaded, it is restricted to a Personal space accessible to only the original uploader and their lab PI. Once approved by the PI, the dataset migrates to the Lab space where others in the same lab will be able to access the dataset. The PI can approve the release of a dataset into the Commons space where other members of the ODC-TBI community will be able to access the dataset. Lastly, the PI can trigger a dataset publication process on the ODC-TBI; once completed, the dataset will be published as a citable unit of research with a unique digital object identifier (DOI) and made accessible to the general public (**Fig 1C**). By granting the PI full control of their data sharing at all times and requiring multiple security checks, we can alleviate security concerns regarding data sharing.

Another common obstacle towards data sharing is the lack of guidance towards adequately organizing the data [31]. Experimental data is commonly stored on spreadsheets with various structures that strive to make the data clearly readable by humans. This includes nested labels, different font sizes, and multiple tables on the same spreadsheet representing different parameters (**S1A Fig**). However, while this approach makes data easy to understand to the original experimenters, the practice creates wide variations in data formats and presents an intractable problem for large-scale data harmonization, interoperability, and merging. The ODC-TBI requires datasets be reformatted into the Tidy format, a standardized data format ideal for data storage, aggregation, and multi-dataset analytics [28] (**S1B Fig**). The ODC-TBI contains written tutorials to guide researchers in formatting their data into the Tidy structure. We also encourage the upload of dataset-associated data dictionaries (**S1C Fig**). Data dictionaries help provide critical definitions for each variable in the dataset, essential information such as the unit of measurement, and additional comments about the experimental protocol that improve the interpretability and reusability of the dataset (e.g. reasons for excluding samples).

To demonstrate the ODC-TBI, we uploaded and aggregated 11 datasets corresponding to 11 prior publications from several labs at the University of California San Francisco [32–42]. Additionally, we included an external dataset from a pooled analysis from the University of Texas Medical Branch published through ODC-TBI and reused under a creative commons (attribution) license (CC-BY 4.0) [43]. The number of animals across all 12 datasets totaled N=1250 individual subjects. The datasets were harmonized according to NIH/NINDS common data elements (CDEs) for preclinical TBI which enabled merging of the datasets for multidataset descriptive analytics as presented in Figure 2. The majority of the uploaded data corresponded to mouse experiments (86.56%). The rest corresponded to rat experiments (13.44%; **Fig 2A**). 74.88% of the subjects were male while 6.24% were female animals. Notably, 18.88% of the records were missing a value for the sex parameter as a result of irrecoverable records which is a common issue when collecting datasets from older publications (**Fig 2B**) [44]. The majority of the experiments utilized the controlled cortical impact contusion injury model (77.6%) with a smaller number of fluid percussion injury (8.56%), closed head injury (6.32%), and closed-head impact model of engineered rotational acceleration (CHIMERA) repeated injury models (7.52%; **Fig 2C**).

**Figure 2.**
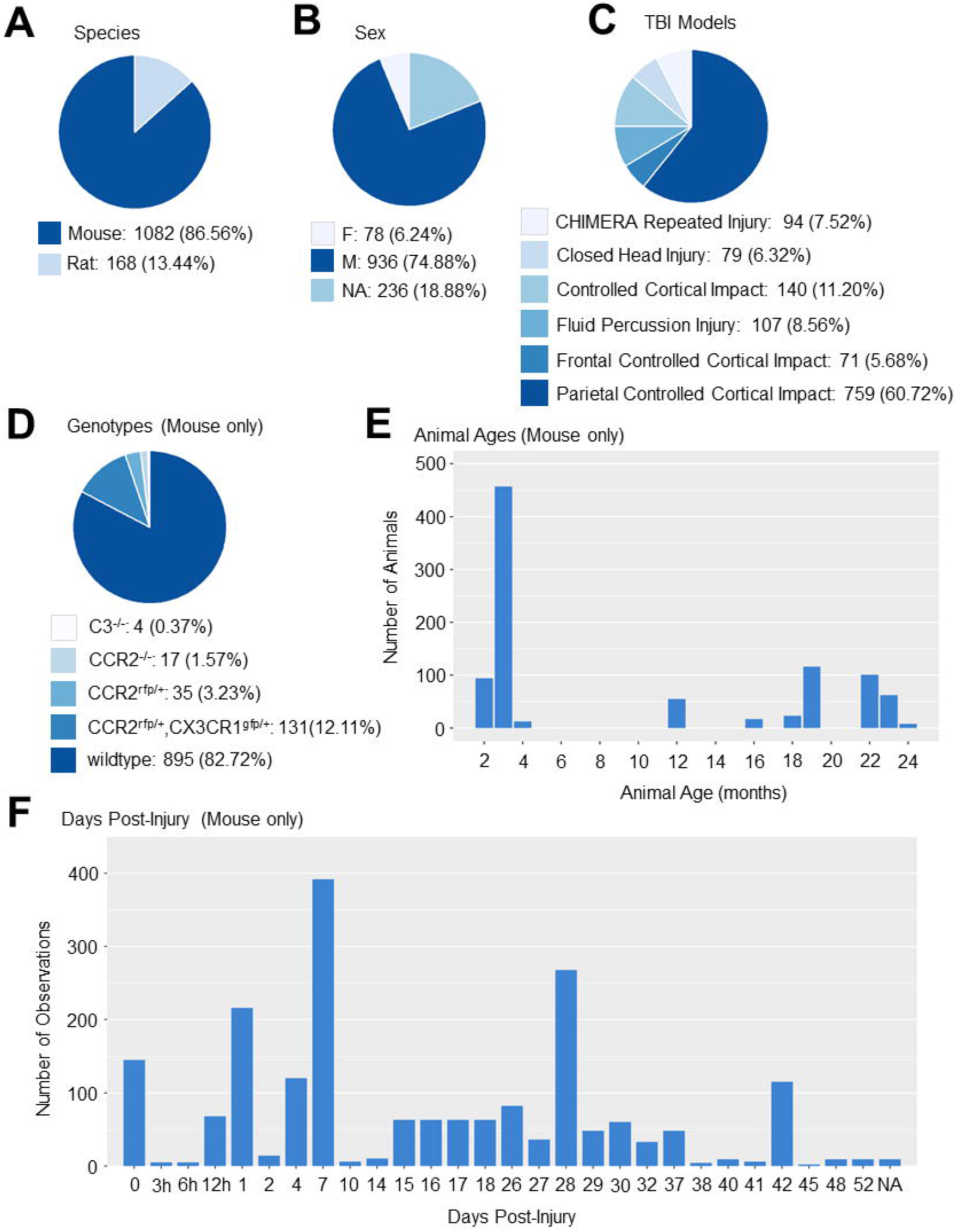
Descriptive summaries of data aggregated from 11 preclinical TBI publications from UCSF on the ODC-TBI. (**A**) The 11 datasets constituted data from 1143 unique animals, 96.66% of which are mice and the remaining 5.34% of which are rats. (**B**) 72.53% of the subjects were male animals and 6.82% were female. 20.65% of the subjects were missing records of male or female. (**C**) The primary TBI model utilized was the controlled cortical impact with 66.4% of experiments utilizing a parietal injury, 6.21% utilizing a frontal injury, and 12.25% not listing the site of CCI in the uploaded dataset. Additionally, 6.91% of the data were from a closed head injury model and 8.22% of the data were from a repeated closed-head injury model utilizing the CHIMERA impactor. (**D**) Of the mice subjects, 82.72% of the subjects were wildtype mice. The remaining subjects included C3-knockout (0.37%), CCR2-knockout (1.57%), CCR2-rfp transgenic (3.23%), and CX3CR1-gfp and CCR2-rfp transgenic (12.11%). These transgenics reflected the interest in inflammatory pathways after TBI in the publications. (**E**) The mice subjects’ age-at-time-of-injury showed a bimodal distribution encompassing young (2-6 mo) and old (16+ mo) animals. The age distribution reflected the focus on the effect of aging on TBI processes. (**F**) Data was collected at a variety of timepoints from the mice experiments. The timepoints with the greatest number of observations were 0 days post-injury (dpi), 1 dpi, 7 dpi, and 28 dpi. The breadth of timepoints reflected time course studies as well as the interest in both acute and chronic effects of TBI in the studies.

We further visualized the characteristics of the 1082 mouse subjects. While most of the mice were wildtype, 17.28% of the subjects were transgenic for immunology-related genes which highlights the fact that the summarized studies were primarily focused on immunological processes of TBI (**Fig 2D**). The subject age distribution showed a bimodal distribution with most animals falling below 6 months or above 18 months of age, reflecting the nature of the studies investigating the effects of age on TBI biology (**Fig 2E**). Lastly, a variety of acute and chronic timepoints were represented in the datasets (**Fig 2F**). Notable peaks in the timepoint distribution included time 0 (often a control timepoint for uninjured animals), 1 day post-injury (dpi), 7 dpi, and 28 dpi to measure acute, sub-chronic, and chronic effects of TBI respectively.

### Missing Data in Data Structure

While working with users to prepare their datasets for upload, we observed that users often had questions regarding uploading files that contain empty cells, also termed “missing values”. Missing values are to be expected: a single dataset can contain data from multiple studies with different outcome measures, resulting in a patchwork of missing and present data. Missing Value Analysis (MVA) is an established statistical subfield that involves descriptive statistical diagnosis of missingness patterns such as whether data is missing completely at random (MCAR), missing at random (MAR), or missing not at random (MNAR or NMAR) [44,45]. Identifying the pattern and reasons for missing data is critical for appropriate data imputation – the statistical practice of replacing missing values with plausible substitute values usually derived from the rest of the data – and multidimensional analytics [44,45]. Dataset-associated methodology and data dictionary documents on the ODC-TBI can be utilized to inform MVA. Here, we highlight common reasons for missingness using the Chou et al 2018 dataset given the authors’ familiarity with the dataset and the breadth of reasons for missingness represented [32]. The simplest visualization for MVA recodes the dataset elements (i.e. spreadsheet cells) with a binary code (0 = missing, 1 = present) and produces a plot of black and grey for missing elements and present elements respectively. MVA revealed that 61% of the elements in the selected dataset are missing values (**Fig 3A**).

**Figure 3.**
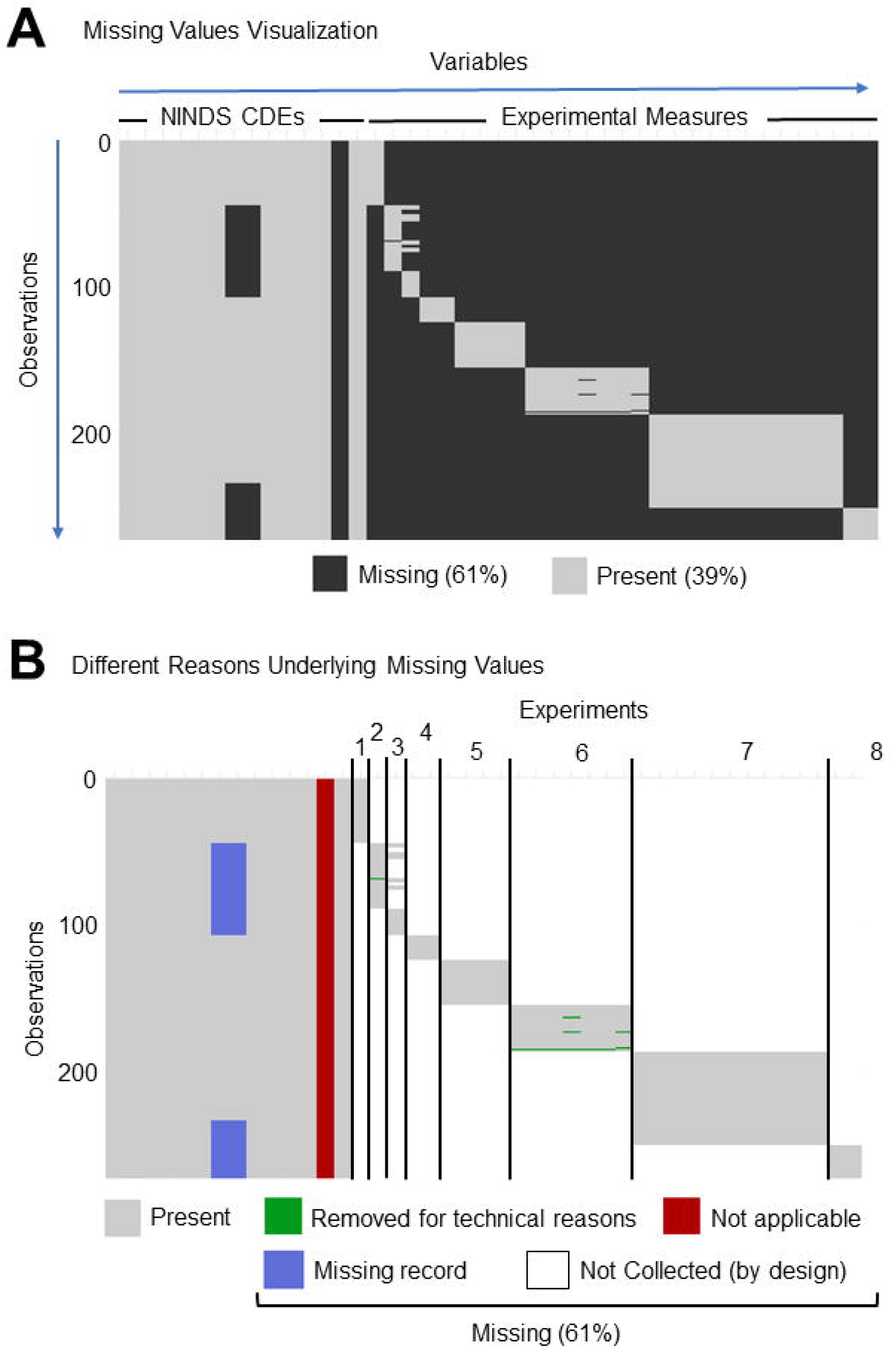
Missing value visualizations of Chou et al 2018. (**A**) Typical missing value visualization shows which elements (i.e. cells) contain a value and which do not, which are thus termed “missing”. The uploaded data showed generally low missingness for variables (i.e. columns) corresponding to NINDS CDEs and fairly high missingness for variables corresponding to collected experimental measures. Each row corresponded to an observation, in this case a single animal subject. (**B**) Types of missingness were manually color-coded based on the type of missingness. The majority of the missing values were “Not Collected (by design)”; the dataset constituted 8 separate experiments, and experimental outcomes were specifically collected for subjects belonging to one experiment. The result was an extremely sparse dataset by design. Another source of missingness was when a variable is “Not applicable,” which we expect in cases when a NINDS-defined CDE is not applicable to the study design. In this example, no treatments were given, so the treatment CDE column was entirely missing values. Data could also be irrecoverable due to “Missing records,” such as the subject’s sex in this example and as reflected in Figure 2B. Lastly, data from experiments could also have been “Removed due to technical reasons”.

Reasons for missingness can be quite varied (**Fig 3B**). Most commonly, measures might not be collected at all as part of the experimental design (“Not collected (by design)” white cells in **Fig 3B**). For example, a sample can be used either for immunohistochemistry or for flow cytometry but not both. Accordingly, two separate cohorts of animals are required: one planned for immunohistochemistry measures and one for flow cytometry. Conversely, there are times when an attempt is made to collect the data, but the data is excluded due to technical reasons (“Removed for technical reasons” green cells in **Fig 3B**). Understanding the circumstances for which the data was removed is critical for the process of data imputation.

In some cases, a variable (i.e. column) may exist in the dataset but not actually be applicable, thus leading to an entire column of missing values (“Not applicable” red cells in **Fig 3B**). This can also be the result of data harmonization and aggregation when certain columns are not applicable to specific data sets. In the Chou et al 2018 dataset, there is a column for the “Treatment” CDE. However, no treatments were administered in any of the experiments, and accordingly the entire column is missing since the parameter was not applicable to the dataset.

Another possible reason for missing values is that the data were not recorded or were unable to be recovered from past records (“Missing record” blue cells in **Fig 3B**). In the Chou et al 2018 dataset, some subjects are missing the sex variable which corresponds with Fig 2B. In this case, the experimental records that we collected the data from did not have the sex information readily available.

### Multidimensional Analytics Use Case

To demonstrate the multi-dataset analytic workflow facilitated by the ODC-TBI, we aggregated data from three controlled cortical impact studies (i.e. independent experimental cohorts of animals) published in Chou et al 2018 and Morganti et al 2015 [32,35]. These studies were chosen because basic multivariate approaches require common variables between datasets. The selected studies included an injury time course study, an aging study, and a treatment study that isolated innate immune cells from injured brain tissue and all measured the expression of the following six inflammatory markers by quantitative PCR: IL-1β, TNF-α, iNOS, Ym1, CD206, and TGF-β. Using the aggregated data, we performed (1) MVA, (2) missing data imputation, (3) principal component analysis (PCA), and (4) syndromic visualization to identify salient multidimensional patterns of immune activation across the studies. After the initial analysis, we included an additional within-study z-score standardization step between MVA and missing data imputation in order to correct for an observed batch effect of study (i.e. study effect) (**Fig 4A**). Broadly, PCA is an unsupervised multivariate technique that combines and reduces the input variables into new features while maximizing the variance of the data accounted for [46,47]. Syndromic visualization encapsulates a set of plots (e.g. syndromic plots, barmaps, heatmaps) developed in the Ferguson lab to intuitively present the PCA results [48,49].

**Figure 4.**
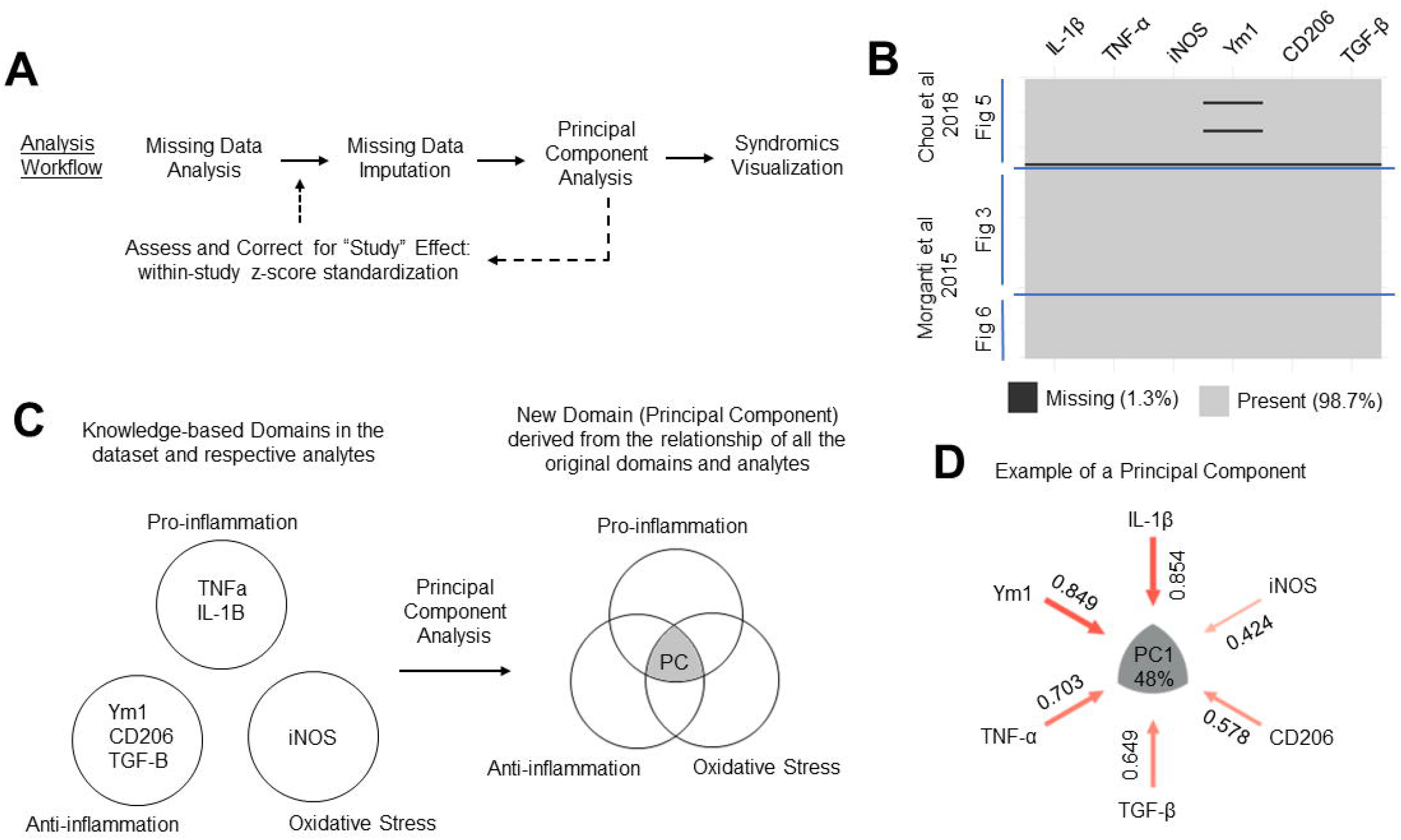
Multidimensional analytics use case. (**A**) We implemented an analysis workflow including missing data analysis, missing data imputation, principal component analysis (PCA), and syndromic visualizations. After an initial analysis, we implemented an additional z-score standardization step prior to data imputation to correct for a study effect. (**B**) Data was aggregated from three experiments (figures) from two papers: Chou et al 2018 and Morganti et al 2015. Visualization of the missing data show 1.3% of the dataset were missing values. Notably, one entire row was entirely missing, and the other two missing values were from the Ym1 variable. (**C**) Conceptual representation of PCA. The original variables (TNF-α, IL-1β, Ym1, CD206, TGF-β, and iNOS) were categorized into the domains of pro-inflammation, antiinflammation, and oxidative stress based off existing knowledge. PCA captures the underlying relationship between the variables – and thus the knowledge domains – to derive new domains from the data. (**D**) The derived PC can be represented as a syndromic plot which visualizes the contributions (i.e. loadings) of each variable to the PC. Furthermore, the PC captures a portion of the variance in the data which is reflected by the percentage value in the center of the syndromic plot. In the example PC, 48% of the variance in the dataset was accounted for, and all six of the variables were loading positively.

MVA revealed that the aggregated dataset has 1.3% missingness (**Fig 4B**). Within the Chou et al 2018 data, one of the samples (i.e. rows which correspond to individual mice in the selected datasets) was missing values across all six variables. A sample that is missing values across all variables cannot be accurately imputed, so we removed the sample from further analysis. Two other samples were missing values for Ym1. Harnessing experimenter knowledge, we identified that the two values are missing due to technical errors during the qPCR procedure. We applied the Little’s statistical test to assess whether the missing data pattern significantly deviated from the null assumption of missing completely at random (MCAR). The result suggested that the missing data were indeed MCAR (p-value = 0.86) [50], indicating that the pattern of missingness was due to random chance and not correlated with the values of other variables in the dataset. Importantly, data imputation could accordingly proceed without the need to explicitly model the missingness [44,45]. We imputed the two missing values using a multiple imputation method that operates under the MCAR assumption: predictive mean matching (PMM). PMM first creates a predictive model from the samples with complete cases (rows without missing values) to generate estimates for the missing values, identifies which complete samples have observed values closest to the predicted value for the missing entry, and then randomly chooses one of the observed values to use for the imputation. We repeated the process ten times – a general guideline for multiple imputation of missing data that is sufficient for cases where only a small portion (< 10%) of the data is missing [51] – to create ten imputed datasets.

We then performed PCA on each individual imputed dataset with mean centering and z-scaling of the data (i.e. analogous to running PCA on the correlation matrix of the dataset) to examine the relationship between the six inflammatory markers. In brief, during experimental design, researchers select outcome measures (i.e. variables) that represent broader domains of interest. In our use case, the variables are inflammatory markers that can be categorized into pro-inflammatory, anti-inflammatory, and oxidative stress domains [32,35]. PCA transforms the variables of this multivariate dataset into a new set of principal components (PCs), also called dimensions, that capture the relationship of the analytes. PCA maximizes the variance in the data along each PC under the restriction that each component is uncorrelated to the others [46,47]. The resulting PCs form data-derived scores determined by patterns in the data (**Fig 4C**). We can further visualize the contributions (i.e. loadings) of each of the original variables to each PC in the form of a syndromic plot (**Fig 4D**).

To determine whether our imputation method significantly affected our PCA output, we tested the similarity of the resultant PCs from PCAs performed on each of the individual PMM-imputed datasets. We found that the resultant PCAs were almost exactly identical (Congruence Coefficient > 0.999 ± 0.001 for each PC; Cattell’s salient similarity = 1 ± <0.001 for each PC). Accordingly, we took the mean of the imputed values to create a single imputed, complete dataset and then applied PCA for further analysis.

The resulting loadings of each variable to each PC allowed us to transform the original data into PC scores for each subject. Plotting the PC scores on the first two PC axes, we observed that the first two components account for variance attributable to study two and study one (along PC1 and PC2 respectively; **Fig 5A**). A two-way analysis of variance (ANOVA) of PC1 scores revealed significant main effects of Injury (F_(1,93)_=8.30, p<0.005) and Study (F_(2,93)_=18.32, p<0.001) and a significant interaction of Injury and Study (F_(2,93)_=4.84, p<0.05). Along PC2, two-way ANOVA similarly revealed significant main effects of Injury (F_(1,93)_=46.69, p<0.001) and Study (F_(2,93)_=4.64, p<0.05) and a significant interaction of Injury and Study (F_(2,93)_=5.63, p < 0.005). To further emphasize the study effect captured by the PC scores, we filtered for all adult sham animals and adult TBI animals at 7 days post-injury across the three cohorts. We observed that the TBI animals fell on both sides of the sham animals along PC1, suggesting that the PCA had transformed the original data according to variance attributable to study as well as biological differences due to injury (**Fig 5B**).

**Figure 5.**
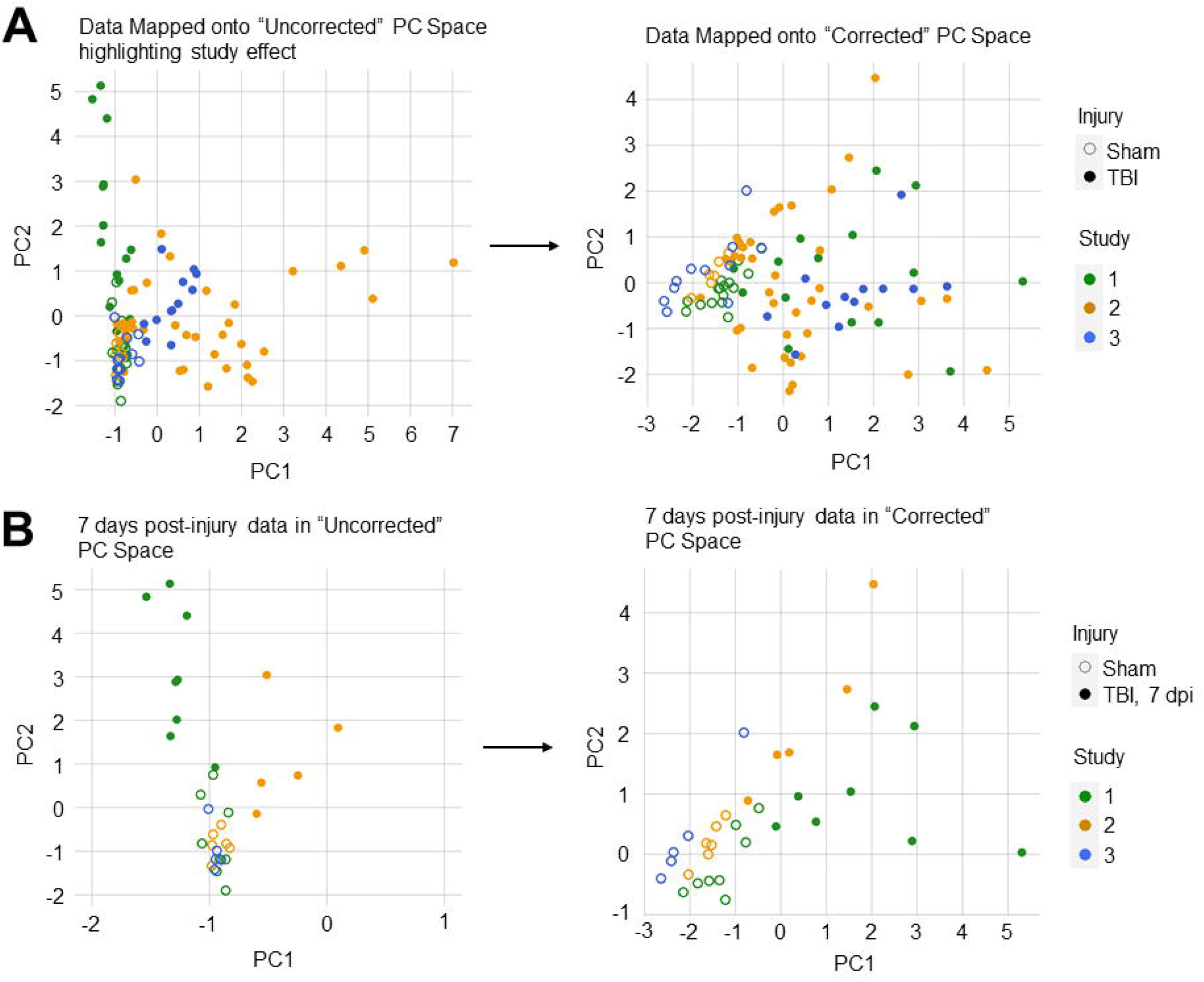
Change in PC scores after correcting for study. (**A**) The datapoints mapped onto PC space (i.e. PC1 and PC2) grouped by Study and Injury groups. In the uncorrected PC space, PC1 primarily captured the variance from Study 2 while PC2 primarily captured the variance from Study 1 (left). Two-way ANOVA revealed significant main effects of Study and Injury and significant interaction along PC1 (Injury main effect: F_(1,93)_=8.30, p<0.005; Study main effect: F_(2,93)_=18.32, p<0.001; Injury and Study interaction: F_(2,93)_=4.84, p<0.05) and PC2 (Injury main effect: F_(1,93)_=46.69, p<0.001; Study main effect: F_(2,93)_=4.64, p<0.05; Injury and Study interaction: F_(2,93)_=5.63, p < 0.005). After correcting for study, PC1 primarily captured the variance between Sham and TBI samples, and neither PC1 nor PC2 appeared to represent the variance from a single study (right). Two-way ANOVA revealed only a significant main effect of Injury along PC1 (F_(1,93)_=84.42, p<0.001). (**B**) The datapoints for animals belonging to similar experimental groups mapped onto the uncorrected and study-corrected PC spaces. Before correcting for the study effect, adult animals at 7 days post-injury (dpi) from study 1 and study 2 fell on opposite sides of the sham experimental groups (left). After correcting for study, the 7 dpi animals clustered more closely in the PC space and exhibited similar PC1 direction in relation to sham animals (right).

To correct for this study effect, we standardized each inflammatory marker into z-scores of the distribution of values within the individual studies (i.e. within-study z-score standardization; **Fig 4A**). After the correction, we reperformed PMM imputation and PCA on the standardized dataset. Two-way ANOVA on the new PC scores revealed only a main effect of Injury along PC1 (F_(1,93)_=84.42, p<0.001). There were no significant main effects or interactions with Study for either PC1 or PC2, suggesting successful correction. Visualization of the study-corrected PC scores also showed that PC1 now primarily captured the variance due to injury (**Fig 5A, 5B**), verifying that our within-study z-score standardization helped correct for the variance between studies that may have been caused by different experimenters.

Taking the PCA of the dataset that had been standardized to within-study z-scores and then averaged across ten imputations, we plotted the variance accounted for (VAF) by each PC on a Scree plot. We observed that the first three PCs (PC1, PC2, PC3) accounted for 83.5% of the variance in the aggregated dataset (**Fig 6A**). We focused our attention on these three PCs given they explain the majority of the data variance and have biologically interpretable loading patterns; conversely, PC4-6 essentially captured unexplained variance and noise in the data. We visualized the loadings of each individual marker to the first three PCs using the syndRomics package in R to generate syndromic plots (**Fig 6B**) [49]. The markers visualized in the syndromic plots were those with absolute loadings above a threshold of significance (|loading| > 0.2). We also visualized the PCA output as a barmap and heatmap which show the loadings for all six inflammatory markers to each PC (**Fig 6C; S2 Fig**). Authors with domain expertise in preclinical TBI neuroinflammation examined the loading patterns and labeled PC1 as representative of an “overall inflammation” axis with every inflammatory marker loading positively and PC2 as representative of the “pro- vs anti-inflammatory” axis with antiinflammatory markers (CD206 and TGF-β) loading inversely to pro-inflammatory markers (IL-1β and TNF-α). Lastly, PC3 showed iNOS loading almost exclusively, suggesting that iNOS might provide unique information about the inflammatory state after injury distinct from the other five markers.

**Figure 6.**
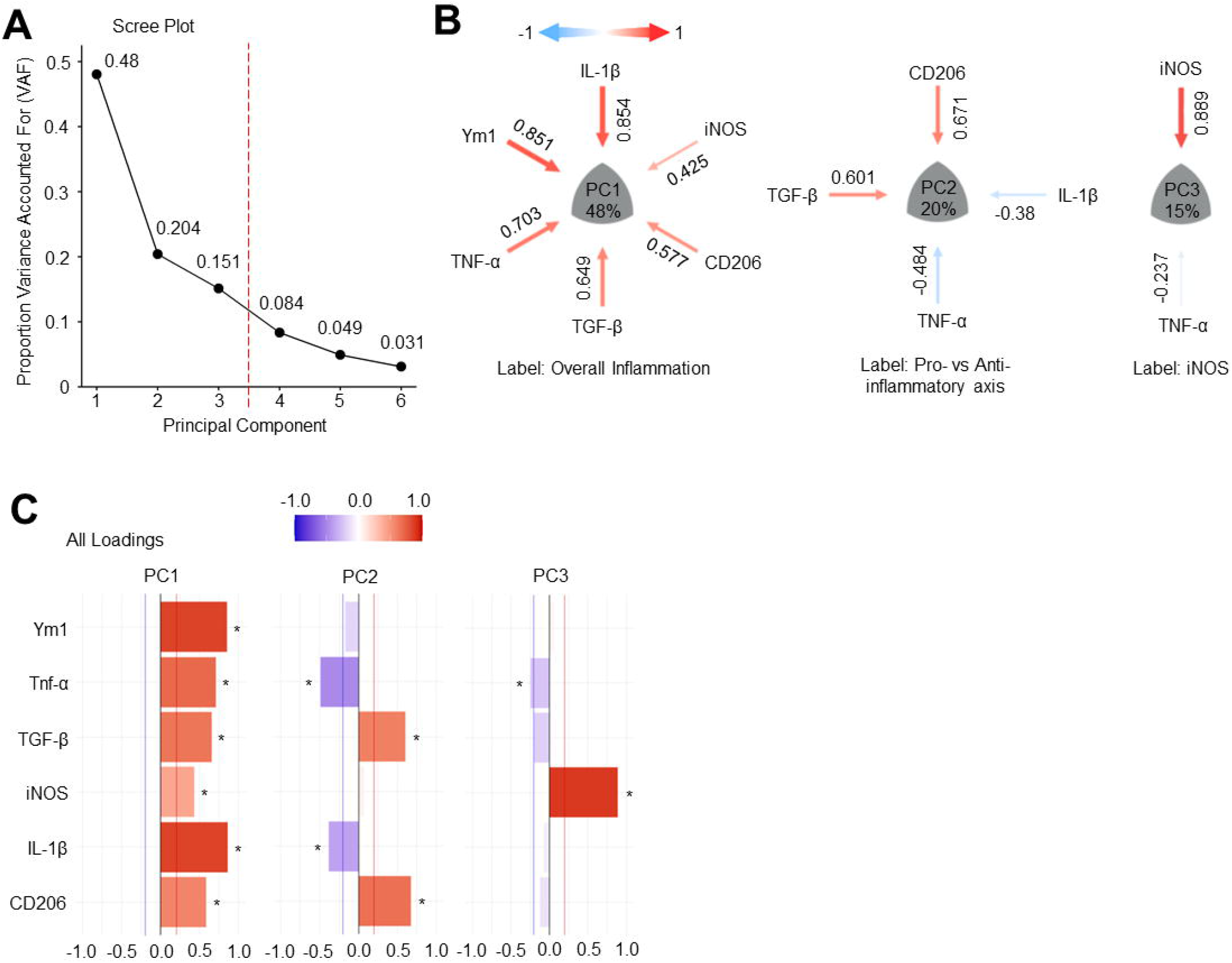
Syndromic visualization of the principal component analysis (PCA). (**A**) The scree plot after running PCA on the imputed dataset revealed that the first three principal components (PCs) account for 83.5% of the variance in the aggregated dataset. (**B**) Syndromic plot visualization showed the significant variable loadings for each PC. PC1 was labeled as overall inflammation, PC2 as the pro- vs anti-inflammatory axis, and PC3 as iNOS expression. (**C**) The barmap visualization provides additional information including the variable loadings that were below the threshold of significance (0.2) for each PC. The barmap denotes which loadings were above the significance threshold with an asterisk.

Notably, data aggregation and PCA can increase the sensitivity (increased effect sizes) to better distinguish experimental groups as compared to univariate analyses. To illustrate this, we mapped the PC scores for adult (3-6mo) and aged (18+ mo) animals in the sham or 7 dpi experimental groups from the aggregated dataset (n=47) (**Fig 7**). We observed that TBI increases the inflammatory profile at 7 dpi as represented by an increase in PC1 score (Injury main effect: F_(1,43)_=65.14, p < 0.001). Furthermore, we observed a distinct separation between adult and aged animals at 7 dpi along PC2: aged TBI animals have lower PC2 scores as compared to adult TBI animals, reflecting an age-driven shift towards pro-inflammation and away from anti-inflammation at the sub-chronic timepoint (two-way ANOVA; Injury main effect: F_(1,43_)=7.44, p<0.01; Age main effect: F_(1,43)_= 18.69, p<0.001; Injury and Age interaction: F(_1,43)_=15.02, p<0.001. Tukey HSD: Adult TBI vs Aged TBI, p<0.001). For comparison, we also reproduced the univariate analyses from Chou et al 2018 with the individual inflammatory markers and the study-specific cohort [32]. We calculated the effect sizes (η^2^) and corresponding observed statistical power (1-β) for the main effects and interactions of Injury and Age for PC1, PC2, and each individual marker (Supplementary Table 1). PC1 had the largest effect size for Injury (η^2^ = 0.593) with an observed power of 1.0. Of particular interest to the original study which examines Injury and Age interactions, PC2 had the largest effect size for the interaction term (η^2^ = 0.179) with an observed power of 0.96. Importantly, the PCs are derived mathematically from correlations in the data and directly model the relationship between variables without relying on multiple univariate comparisons which would be prone to false positives. PCA thus not only improved sensitivity for the original experimental question but also specifically established that age skews sub-chronic inflammation away from anti-inflammation and towards pro-inflammation. This was not clearly observed in the original cohort and univariate analyses with the individual markers.

**Figure 7.**
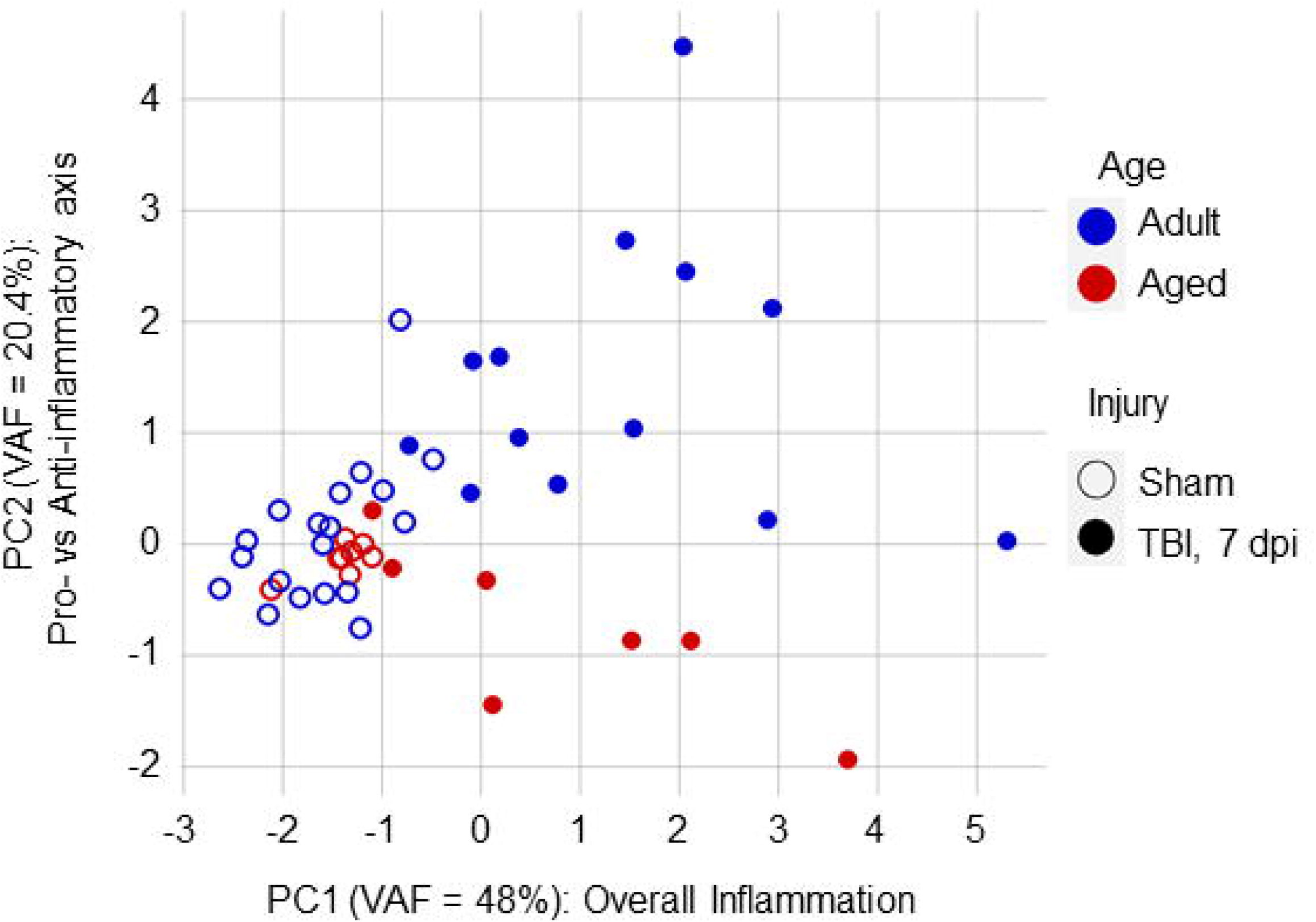
Validation of results from previous studies with the aggregate analysis. Animals corresponding to sham vs TBI at 7 days post-injury and Adult (3mo) vs Aged (18+ mo) experimental groups were filtered from the aggregated dataset and mapped onto the study-corrected PC space. Sham animals clustered closely regardless of age. TBI increased the overall inflammation (PC1) for TBI animals (two-way ANOVA; Injury main effect: F_(1,43)_=65.14, p < 0.001) without a significant main effect of Age or interaction. Along PC2, aged animals exhibited a shift towards pro-inflammation while adult animals shifted towards anti-inflammation at 7 days post-injury (two-way ANOVA; Injury main effect: F_(1,43)_=7.44, p<0.01; Age main effect: F_(1,43)_=18.69, p<0.001; Injury and Age interaction: F_(1,43)_=15.02, p<0.001. Tukey HSD: Adult TBI vs Aged TBI, p<0.001).

Altogether, our analyses demonstrate the potential of multidimensional analytics in reinforcing inferential reproducibility of previous findings while leveraging heterogeneous datasets to identify novel pathophysiological patterns. By establishing a functional infrastructure towards the FAIR principles of data sharing to promote data reuse, the ODC-TBI acts as a critical bridge for dataset standardization and aggregation to facilitate and accelerate such efforts.

## Discussion

The ODC-TBI is a data commons developed for preclinical TBI research designed to (1) enable data sharing within the research community [12], (2) support data standardization guidelines established by the NINDS [20,21], (3) promote FAIR data sharing principles [25], and (4) empower Big Data analytics in preclinical TBI research [10]. Through the ODC-TBI, we leverage the heterogeneity of preclinical TBI research to identify common TBI immune responses across three different preclinical TBI studies that vary time post-injury, age, and treatment. Our multivariate, multidimensional analysis reveals patterns of inflammation that corroborate the current paradigm of a pro- and anti-inflammatory axis as well as highlight the unique relationship of oxidative stress to other inflammatory markers. The ODC-TBI will enable researchers to identify common patterns of TBI pathophysiology persistent across injury models, injury parameters, experimental designs, and different research labs themselves by harnessing data at the level of the individual subject.

We recognize that the practice of data sharing is still emerging in many biological fields including preclinical TBI. There are perceived risks of data sharing that endure among research communities even as publishers and funding agencies have begun to require it [23,31,52]. The issues of data security, the possibility of being scooped, or the chance of the data being misused or used without proper citation are all common concerns highlighted by several survey studies [29,30]. With the ODC-TBI, we ensure that the PI has full control of the accessibility of their dataset when sharing their work with their peers in the research community and when they publish the datasets to the general public. Additionally, ODC-TBI also tracks which users have accessed shared datasets and provides the information to the dataset PIs. During dataset publication, PIs provide critical metadata such as dataset authors and contributors, and the ODC-TBI generates a unique and persistent digital object identifier (DOI) and citation for the dataset. Most importantly, all datasets published through the ODC-TBI are done so under the Creative Commons CC-BY 4.0 license, meaning any work utilizing those datasets must properly cite them much the same way scientific papers are cited. This provides a novel avenue for researchers to benefit from their data as a new citable scientific work product. This has direct benefits to the data contributor, as data sharing has been found to be associated with an increase in citations for researchers [53]. These requirements are explicitly written as part of the data use agreement consented to by all users signing up to the ODC-TBI and provide a key layer of accountability. Future features of the ODC-TBI platform will include direct peer-to-peer sharing functionalities, further diversifying the methods PIs can upload and share their data in a protected manner on ODC-TBI.

We also realize the importance of supporting and integrating common terminology such as CDEs to improve all aspects of FAIR data sharing on the ODC-TBI. Indeed, NINDS and the TBI research community have recognized the challenges in data comparison due to the lack of common variable names and definitions; this spurred a concerted multi-center endeavor to identify and define CDEs to be adopted by clinical and preclinical TBI researchers [15,20,21]. We manually aligned variables in 11 datasets described here to NINDS-defined CDEs prior to uploading them to the ODC-TBI. To promote the practice of aligning NINDS-defined CDEs, we aim to implement a CDE mapping system on the ODC-TBI built upon the engineering framework of the InterLex/NeuroLex system developed by NIF [22]. The feature will enable CDE mapping after data upload and allow users to align each variable of a dataset to a dictionary of CDEs (including NINDS-defined CDEs). The mapping system would increase the accessibility and prominence of the NINDS-defined CDEs and help construct a knowledge base of TBI research to make data more findable, interoperable, and reusable. As the number of datasets shared on the ODC-TBI grows, it will be possible to further validate the prevalence of NINDS-defined CDEs as well as identify novel CDEs in TBI research.

We expect many datasets uploaded to the ODC-TBI to contain missing values for a variety of reasons such as those visualized in Figure 3. MVA is a critical component for Big Data analytics; many multivariate techniques as well as common univariate approaches (t-test, correlation, ANOVA) require complete datasets for analysis. Most commercial statistics tools default to dropping subjects (listwise deletion) with missing values. However, researchers are often unaware of the impact of missing values, and the practice of listwise deletion can introduce bias and contribute to scientific irreproducibility [44,50]. Understanding the types of missingness is essential for selecting which data imputation technique can be applied; various imputation techniques contain different assumptions that would invalidate specific analyses if they are violated [44,54]. In simple cases such as when the Treatment column is not applicable in the study but still kept as a column (red-labeled cells in **Fig 3**), the missing value can be imputed with a control value (e.g. control, naïve, 0). More generally, data imputation depends on modeling the correlation between variables, and if two variables are never collected in tandem because of experimental design or limitations, then identifying their relationship becomes increasingly inaccurate. Recognizing when data is missing because of experimental design is critical. To this end, the ODC-TBI supports the upload of dataset-associated methodology and data dictionaries that can provide context for researchers to interpret when data imputation is appropriate.

Similarly, the methodology documents and data dictionaries can also highlight the reasons data may be missing due to technical reasons. In many cases, data is missing because of truly random events (e.g. contamination of a single sample during processing or human error performing the experimental protocol). In such cases, the missing data would be classified as missing completely at random (MCAR), which permits straightforward approaches to imputing missing data without having to incorporate patterns of missingness directly into the analysis [44]. In other circumstances, the data might be missing not at random (MNAR): the missingness is correlated to one of the other variables of interest or itself. An example of MNAR data is when a sample’s protein quantification falls below the detectable range of an assay. Instead of keeping a potentially inaccurate value, the experimenter decides to exclude the value altogether. Here, the missingness is correlated to the variable itself: the value is missing because its value falls below the detectable limit. If we were to impute the missing values without regard, we would overestimate the true values of the missing data and bias our analyses. With proper documentation, such cases of MCAR, MNAR, and missing at random (MAR) data can be identified to better inform the appropriate approach to data imputation and avoid grave statistical mistakes in analysis.

As an illustration of how ODC-TBI data can be reused for further discovery, we pooled data across three cohorts of subjects from prior papers [32,35] and performed multivariate analysis. In the original univariate analysis of Chou et al 2018, we observed a general impairment of anti-inflammatory markers but no difference in pro-inflammatory markers at 7 days post-injury in aged animals after TBI [32]. While the univariate analysis suggested a shift in pro- vs anti-inflammatory states, at best we could conclude that the anti-inflammatory response after TBI was blunted by age. Our multivariate analysis shows that across the three studies, there is a distinct PC (PC2) that can be interpreted as the pro- vs anti-inflammatory state as measured by CD206, TGF-β, IL-1β, and TNF-α. From PC2, we were able to expand on the original results and infer that age does bias sub-chronic inflammation after TBI towards proinflammation. Additionally, while Ym1 is considered an anti-inflammatory marker on myeloid cells [55], our results suggest that Ym1 is not as informative as CD206, TGF-β, IL-1β, and TNF-α in explaining the pro- vs anti-inflammatory state of the tissue. Ym1 could be excluded from measurement for studies primarily focused on determining the pro- vs anti-inflammatory state. While myeloid cells do not exhibit strictly pro- or anti-inflammatory phenotypes after TBI [35,36], our analyses show that in the aggregated dataset, there is a marked inverse relationship between pro-inflammatory (IL-1β, TNF-α) and anti-inflammatory markers (CD206, TGF-β). Overall, the PCA corroborated and augmented previously published research [32,35] while increasing the statistical power and inferential validity by analyzing a larger sample (N = 99) through dataset aggregation and directly modeling the underlying patterns with a multidimensional approach.

Additionally, our analysis isolates the expression of iNOS in PC3. While expression of iNOS has been correlated with pro-inflammatory responses in innate immune cells after TBI [55,56], oxidative stress is largely considered a related-but-separate secondary injury mechanism [57]. Our results reinforce this perspective with innate immune cells because of the PCA property that the resultant PCs are uncorrelated. Correspondingly, the PCA supports (1) modulating inflammation and oxidative stress as two complementary therapeutic approaches and (2) targeting innate immune cells as an intersection of both. Importantly, the data we analyzed were obtained specifically from isolated innate immune cells and thus do not capture the relationships with other secondary injury mechanisms or other oxidative stress pathways. As more datasets are shared and published, further multivariate analyses can be done to associate or dissociate secondary injuries more holistically and with greater sensitivity.

Notably, the analytical workflow presented here can be extended to reveal features persistent across laboratories, experimenters, injury parameters, and injury models. This approach is powerful because it ultimately leverages the heterogeneity of experimental design in preclinical TBI research to find common underlying pathophysiology of TBI. Critically, while the PCs are extracted through multivariate statistical techniques, assigning labels and contextualizing the PCs with the underlying biology requires a combination of statistical rules based in the well-established field of factor analysis [58,59] as well as the specific biomedical domain expertise. Data sharing through the ODC-TBI will open avenues of collaboration not only between researchers in preclinical TBI but also between computational and molecular researchers who can provide complementary expertise towards interpreting results. As more studies populate the ODC-TBI, such opportunities and interdisciplinary collaboration will identify features of TBI across an even broader array of heterogeneity and uncover possible therapeutic targets and biomarkers that would be applicable to a broader patient population.

## Materials and Methods

### Data Formatting and Upload to the ODC-TBI

Data from 11 published studies at UCSF were collected from various data sources and structured according to the Tidy data format [28]. Variable names were aligned to NINDS preclinical TBI CDEs when possible [20,21]. Data was uploaded following the ODC-TBI data upload workflow. The specific variables and data analyzed in this paper will be published and made accessible on the ODC-TBI.

### Data Summarization and Missing Values Visualization

The datasets were downloaded from the ODC-TBI and aggregated using the open-source programming language R [60]. Data summaries were generated using *tidyverse* [61] for dataframe manipulation and *ggplot2* [62], *RColorBrewer* [63], and *colorRamps* [64] R packages for visualization.

Missing values visualizations were generated using the “vis_miss” function in the *naniar* R package [65]. Labeling of the types of missingness was done manually by relying on researcher familiarity with the dataset.

### Multidimensional Use Case Workflow

Quantitative PCR measures of six cytokines (IL-1β, TNF-α, iNOS, Ym1, CD206, and TGF-β) from three experiment cohorts [32,35] were combined into a single dataset. A missing values visualization was generated using the *naniar* R package, and rows that were missing values across all cytokine variables (i.e. columns) were removed (1 row removed). Little’s missing completely at random (MCAR) test was performed using the “LittleMCAR” function in the *BaylorEdPsych* R package [66] to determine whether the pattern of missingness is not MCAR (i.e. missing at random (MAR) or missing not at random (MNAR)) [50].

To impute missing values, we used the “mice” function in the *mice* R package [67] with the parameters: 10 imputations, predictive mean matching method, and a seed value of 200. Linear Principal Component Analysis (PCA) was performed using the “prcomp” function in the *stats* R package [60] with centering and scaling [68,69].

To correct for the batch effect (i.e. due to study), we added a z-score standardization step after removing the rows missing data across all columns and prior to data imputation. We first calculated the mean and standard deviation for each cytokine for each of the three studies. For each cytokine datapoint, we then subtracted the respective mean and divided by the standard deviation of the study.

To determine the stability of the PCA results across the 10 imputations generated through *mice*, we utilized the “component_similarity” function in the *syndRomics* R package [49]. We reported the resulting Congruence Coefficient and Cattell’s salient similarity metrics. Because the PCA outputs for each imputation were highly similar, we averaged all 10 imputations together to generate the final imputed dataset. PCA was then performed on the imputed, averaged dataset for further analysis.

To visualize the scree plot, we calculated the variance accounted for (VAF) for each principal component (PC) from the “sdev” output of “prcomp”:

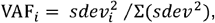

where *i* is the PC number and the denominator is the sum of the variance across all PCs. We selected the top PCs that collectively explained over 80% of the variance in the data and had biological interpretations. PC loadings were calculated and visualized using the “syndromic_plot”, “barmap_loading”, and “heatmap_loading” functions in the *syndRomics* R package. PC scores were obtained from the “x” output of “prcomp” which transformed the original variables into values along each PC.

To determine the study and injury effects (**Fig 5**) and to determine the injury and age effects (**Fig 7**), we performed two-way ANOVA with Tukey HSD posthoc using the “aov” and “TukeyHSD” functions in the *stats* R package respectively.

To compare effect sizes and observed power, we performed two-way ANOVA for the main effects and interaction of Injury and Age on PC1 and PC2 of adult and aged sham animals and animals at 7 days post-injury from the aggregated dataset (n = 47). We additionally filtered for the Chou et al 2018 cohort (n = 31) and performed two-way ANOVA on the six individual inflammatory markers. The effect size (η^2^) of the Injury effect, Age effect, or interaction was calculated from the ANOVA F table as:

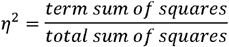

To obtain observed power, we calculated the partial η^2^:

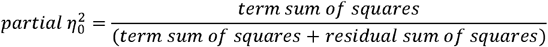

We then converted the partial η^2^_0_ (which is based on sample estimates) to the partial η^2^ based on Cohen’s f according to the G*Power manual [70]:

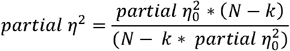

where N is the total number of samples and k is the total number of groups in the experimental design. Partial η^2^ was converted to Cohen’s f:

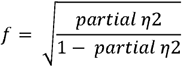

The observed power was then calculated from Cohen’s f using the “pwr.f2.test” function in the *pwr* R package [71].

---

**SFigure 1.**
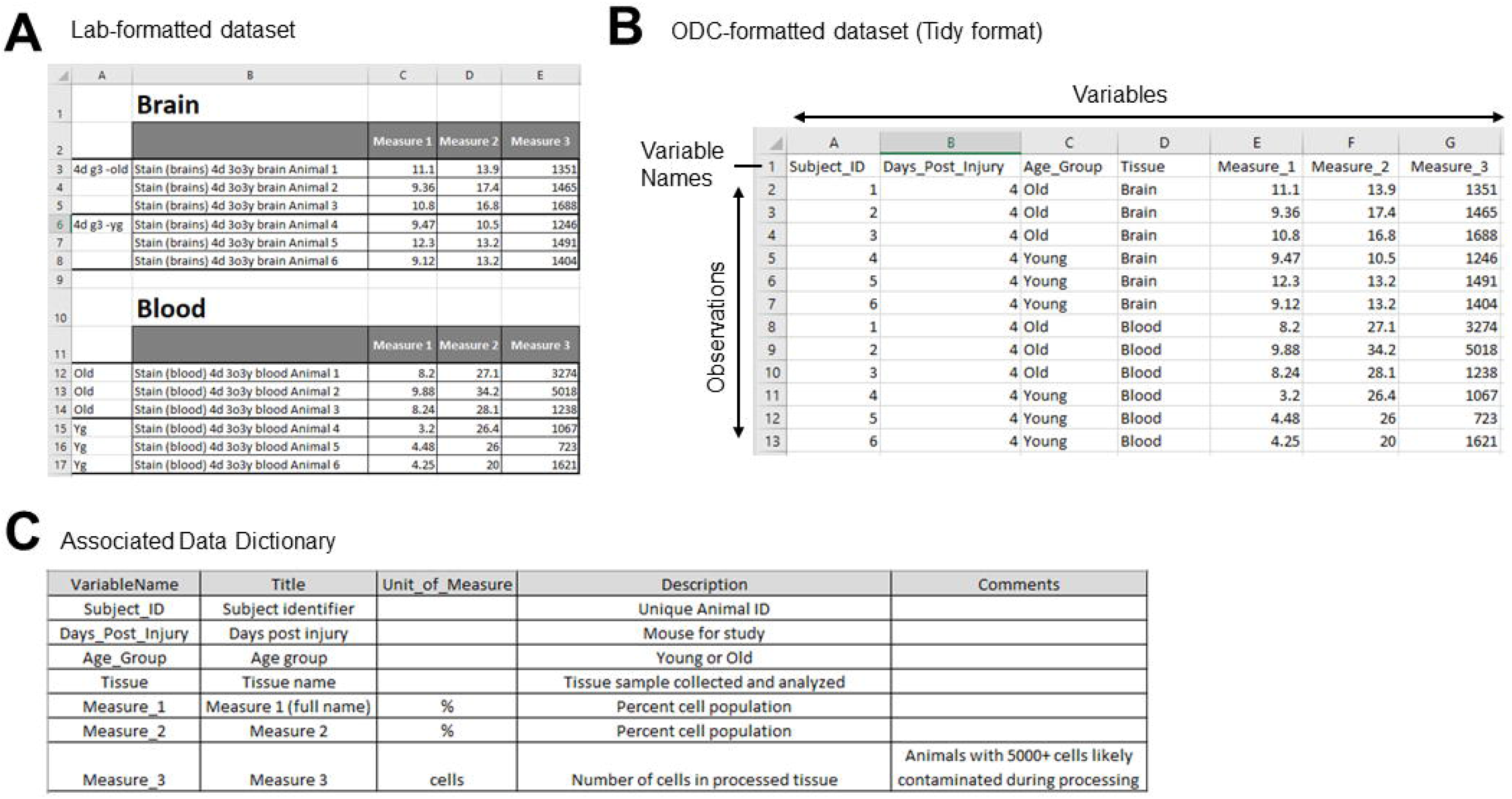
Data and data dictionary formatting for ODC-TBI upload. (**A**) Experimental data is commonly recorded in spreadsheets with various structures designed to be human-readable including multiple tables and nested labels on a single spreadsheet. (**B**) ODC-TBI requires data to be formatted into the Tidy format. The first row contains the variable (i.e. column) names, and each column represents one of the dataset variables. Each corresponding row contains the values for an observation. (**C**) ODC-TBI allows the upload of a data dictionary with each dataset. The ODC-TBI data dictionary contains the following five columns: VariableName, Title, Unit_of_Measure, Description, and Comments.

**SFigure 2.**
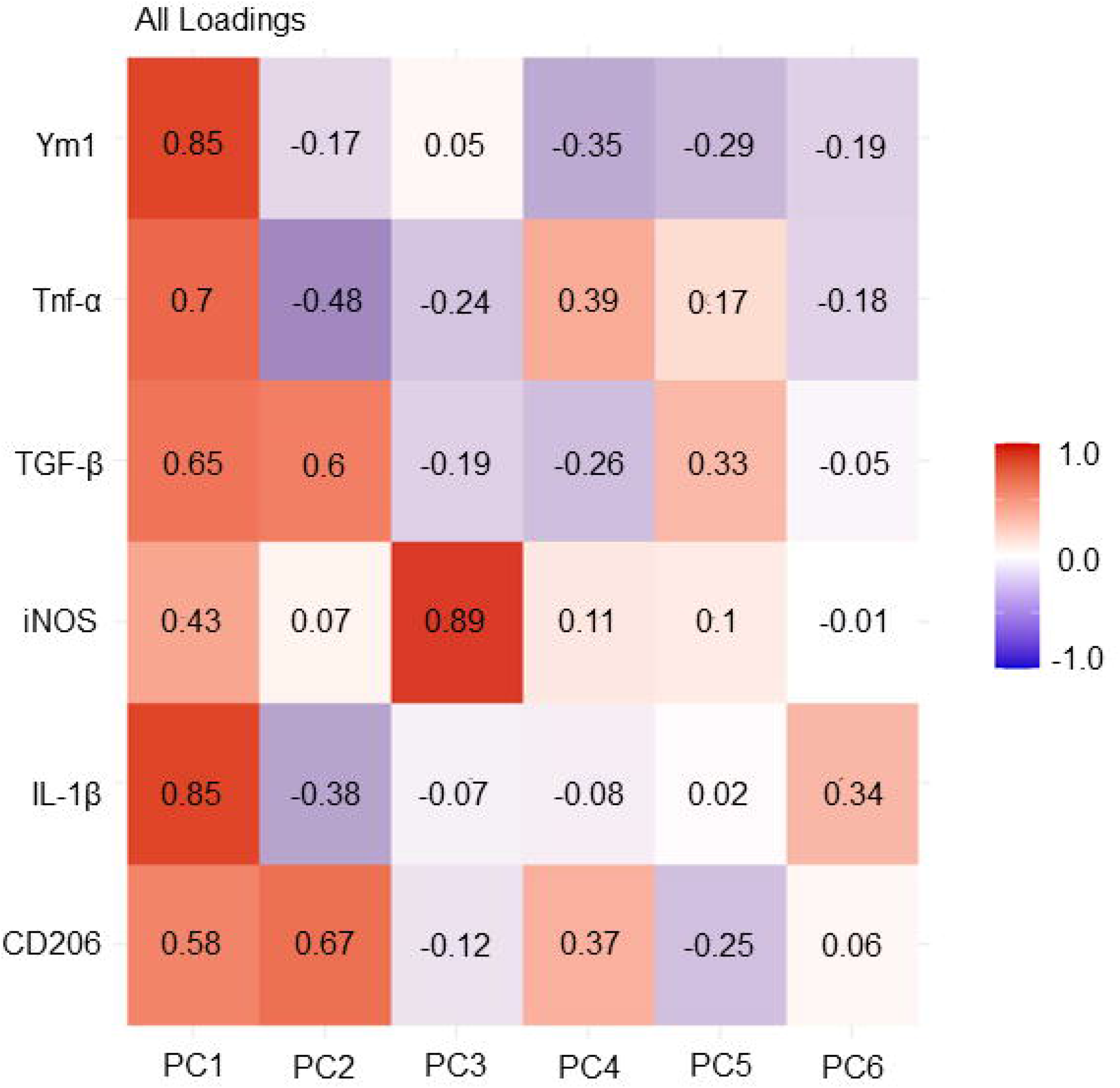
Heatmap representation of PCA. The heatmap visualization provides similar information as the barmap and shows all variable loadings including those below the threshold of significance (0.2) for each PC. The heatmap also shows the loadings for all PCs, including PC4, PC5, and PC6.

**Supplementary Table 1.** Effect size and observed power for Injury and Age effects for PCs and univariate inflammatory markers (sorted by descending effect size for each effect).

| Effect | Variable | Effect Size ($\eta^2$) | Power |
| --- | --- | --- | --- |
| Injury | **PC1** | **0.593513** | **1** |
|  | TGFB | 0.587202 | 1 |
|  | TNFa | 0.528806 | 0.999555 |
|  | IL1B | 0.38067 | 0.968824 |
|  | Ym1 | 0.37861 | 0.982232 |
|  | CD206 | 0.356279 | 0.999821 |
|  | PC2 | 0.088426 | 0.741933 |
|  | iNOS | 0.002289 | 0.05672 |
| Age | CD206 | 0.224005 | 0.992383 |
|  | **PC2** | **0.222114** | **0.98516** |
|  | TGFB | 0.084818 | 0.789029 |
|  | Ym1 | 0.066147 | 0.397578 |
|  | iNOS | 0.051901 | 0.210149 |
|  | TNFa | 0.00562 | 0.084759 |
|  | PC1 | 0.002792 | 0.082704 |
|  | IL1B | 0.001879 | 0.058333 |
| Injury:Age | **PC2** | **0.178517** | **0.959657** |
|  | CD206 | 0.146892 | 0.944496 |
|  | TGFB | 0.067209 | 0.691436 |
|  | Ym1 | 0.062637 | 0.380091 |
|  | iNOS | 0.024693 | 0.124739 |
|  | TNFa | 0.021873 | 0.189681 |
|  | PC1 | 0.011898 | 0.194191 |
|  | IL1B | 0.006931 | 0.081102 |

## Notes

https://odc-tbi.org//

### Competing Interest Statement

The authors have declared no competing interest.

